# Sequential design of single-cell experiments to identify discrete stochastic models for gene expression

**DOI:** 10.1101/2024.09.12.612709

**Authors:** Joshua Cook, Eric Ron, Dmitri Svetlov, Luis U. Aguilera, Brian Munsky

## Abstract

Control of gene regulation requires quantitatively accurate predictions of heterogeneous cellular responses. When inferred from single-cell experiments, discrete stochastic models can enable such predictions, but such experiments are highly adjustable, allowing for almost infinitely many potential designs (e.g., at different induction levels, for different measurement times, or considering different observed biological species). Not all experiments are equally informative, experiments are time-consuming or expensive to perform, and research begins with limited prior information with which to construct models. To address these concerns, we developed a sequential experiment design strategy that starts with simple preliminary experiments and then integrates chemical master equations to compute the likelihood of single-cell data, a Bayesian inference procedure to sample posterior parameter distributions, and a finite state projection based Fisher information matrix to estimate the expected information for different designs for subsequent experiments. Using simulated then real single-cell data, we determined practical working principles to reduce the overall number of experiments needed to achieve predictive, quantitative understanding of single-cell responses.

## 1 INTRODUCTION

Single-cell gene expression can be highly variable due to the inherently discrete number of important molecules, especially genes and RNA [1]. Many genes are only present in as few as one or two copies per cell, and these are activated or deactivated by cellular stimuli to produce large relative fluctuations (over time, or from cell to cell) as these genes switch between transcriptionally active and silent states [2]. This variability in activity is measurable using modern optical microscopy experiments, where specific bio-molecules are labeled and then imaged at sub-cellular resolution. For example, in immunocytochemistry (ICC), fluorescently labeled antibodies are frequently used to quantify the position and intensity of different proteins, or in single-molecule fluorescence in situ hybridization (smFISH), RNA molecules are labeled and counted within within individual cells.

These microscopy experiments quantify the behaviors of cellular processes and, when combined with computational models, can provide greater understanding for gene regulation mechanisms [3]. With single-cell data, one can apply the framework of the chemical master equation (CME) to capture and predict the full distributions of cellular fluctuations [4, 5]. There are a massive number of different single-cell microscopy experiments that one could perform to characterize a cellular response, but these experiments are costly and time-consuming, and the community needs new experiment methods to design which experiments to conduct, how much data to collect, and under what conditions.

The Fisher information matrix (FIM) is a powerful tool that has been used in recent years to assess identifiability and robustness of deterministic ordinary differential equation (ODE) models [6], and in crafting efficient bulk measurement experiments to reduce this uncertainty [7, 8]. FIM has also been used to design experiments for stochastic processes using the linear noise approximations or other statistical moment closure techniques [9, 10] to investigate the impact of various measurement types on parameter uncertainties in stochastic gene expression modeling. More recently, a handful of studies have applied FIM calculations directly to the CME description of chemical kinetics [11, 12] to examine the effectiveness of single-cell experiments with time-varying inputs to identify stochastic models with nonlinear reaction rates and more complex multi-modal single-cell expression distributions, including in situations where measurements are distorted by probabilistic effects such as labeling inefficiencies or image-processing errors [13]. These previous CME-based FIM analyses have demonstrated through simulation and experimental data that the FIM can correctly estimate model uncertainty after models have been fit to sufficient data. However, it remains to be seen how well these tools can perform when prior information or data are limited, or when model mechanisms are unknown.

To address this challenge, we explore a sequential experiment design (SED) strategy that approximates the CME (Sec. 2.1) using Finite State Projections (FSP, 2.2) to compute the likelihood of single-cell data (2.3) and then sample the Bayesian posterior (2.4). We then compute a FSP-based FIM to estimate the expected information (2.6) and select subsequent experiments (2.8). We demonstrate the approach on two simple models with simulated data (III-A-B) and on experimental single-cell ICC data (III-C). We then conclude with practical working principles to reduce the number of experiments needed to achieve predictive, quantitative understanding of single-cell responses.

## 2 METHODS

### 2.1 Chemical Master Equation (CME)

Consider an *N*-species (bio)chemical process with integer population state vector, **x**_*j*_ ≡ [*x*_1_, …, *x*_*N*_]^*T*^ ∈ ℕ^*N*^. This vector can change randomly in time via *M* discrete reactions, **x**_*j*_ → **x**_*i*_ = **x**_*j*_ + ***ν***_*µ*_, where the stoichiometry ***ν***_*µ*_ defines the change due to the *µ*^th^ reaction. Furthermore, define the finite-valued propensity function *w*_*µ*_(**x**, *t*, ***θ***) such that *w*_*µ*_(**x**, *t*, ***θ***)*dt* denotes the probability that the *µ*^th^ reaction will occur in the infinitesimal time step, (*t, t* + *dt*). We assume that *w*_*µ*_(**x**, *t*, ***θ***) is non-negative for **x** ≥ 0 and piece-wise differentiable with respect to *t* and parameters in ***θ*** ≥ 0.

For any enumeration of all possible states, 𝒳 = [**x**_1_, **x**_2_, …], let the probability mass vector be written as **P** = [*P*(**x**_1_|*t*, ***θ***), *P*(**x**_2_|*t*, ***θ***), …]^*T*^. For simplicity, we will drop the conditions (*t*, ***θ***) from the notation and write simply *w*_*µ*_(**x**_*i*_) and *P*(**x**_*i*_). With this notation, the probability of each state evolves according to the CME as:

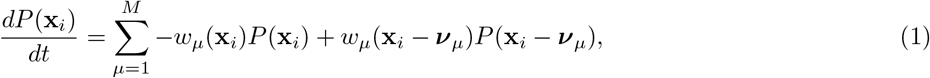

which can be collected in matrix form as:

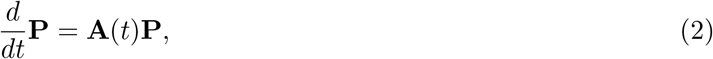

where elements of the infinitesimal generator matrix are:

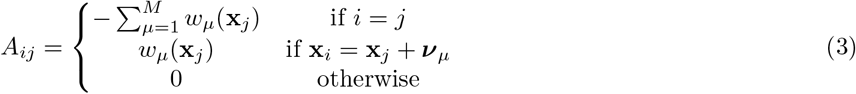

Because the CME contains one ODE for each potential state of the system, its dimension can be infinite, and its exact time-varying solution is unknown except for a handful of simple examples, requiring approximations.

### 2.2 Finite State Projection (FSP) bounds on CME Solution

In the FSP approximation [14] to the CME solution, we choose a finite, ordered subset of *K* states given by 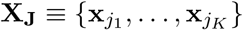. We construct a new Markov process retaining states within this set, but replacing the rest with absorbing sinks. The new master equation becomes:

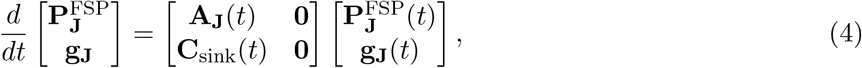

where **A**_**J**_ is the principal sub-matrix of **A** corresponding to the indices in **J**, and the non-negative vector **g**_**J**_(*t*) is the cumulative probability of escape into the specified sinks. The non-negative matrix [**C**_sink_]_*lj*_ quantifies the transition rates from **x**_*i*_ ∈ **X**_*J*_ into the sink *g*_*l*_.

The FSP provides three important guarantees [14]:

1. the FSP solution is a lower bound on the exact solution,

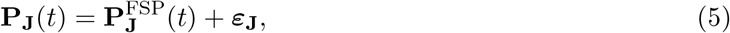

where |***ε***_**J**_(*t*)|_1_ ≤ |**g**_**J**_(*t*)|_1_;
2. the exact error of the FSP approximation over all of the original states, **X**_**J**_ **⋃X**_**J**_*′* = **X**, is calculated as

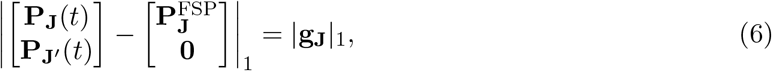
3. the difference between the FSP solution and the true solution decreases monotonically as additional states are added to **J**, such that if **J**_2_ ⊇ **J**_1_, then 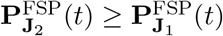 and 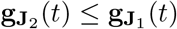 for all *t*.

### 2.3 FSP Bounds on Likelihood Function

Suppose that *d* = {1, 2, …, *D*} *independent* cells have been measured (e.g., using ICC or smFISH) at different times *t*_*d*_, and each measurement **y**_*d*_ corresponds either to a specific state, **y**_*d*_ = **x**_*d*_ or marginalization over multiple states, 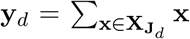. Given a CME model above, and a data set 𝒟 = {**y**_*d*_}, the log-likelihood to make all of these observations given model parameters ***θ*** is:

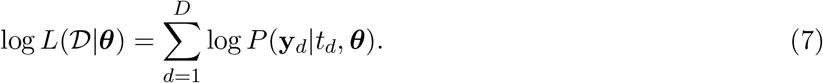

From (5) and (6), we can specify lower and upper bounds on this likelihood function according to:

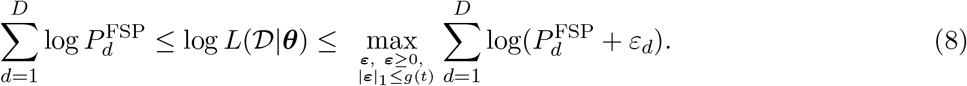

The right-most term can be solved efficiently using a water-filling algorithm [15], and the FSP guarantees that the bounds converge monotonically toward the exact solution as new states are added to **J** (the rate of this convergences depends on the model and the method by which states are added).

### 2.4 Metropolis-Hastings Algorithm to sample posteriors

From Bayes’ law, the posterior distribution given data 𝒟 can be calculated (up to an integration constant) in terms of the prior log-probability distribution log *P*(***θ***) and the log-likelihood function according to:

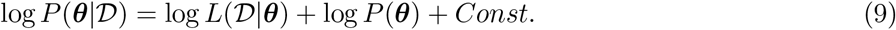

Given the bounds on the likelihood function in (8), one can apply the Metropolis-Hastings algorithm (MHA) to sample from the posterior distribution [16]. For the MHA, parameters are defined in log-space, and we use a symmetric proposal distribution with a covariance equal to the inverse Fisher information matrix (see below).

### 2.5 Sensitivity of the CME Solution

Define the CME *sensitivity* to the *n*^th^ parameter as **s**_*n*_ = *d***P**/*dθ*_*n*_. To our knowledge, proving direct FSP bounds on the sensitivity is an open question. However, one can approximate the sensitivity using finite difference as

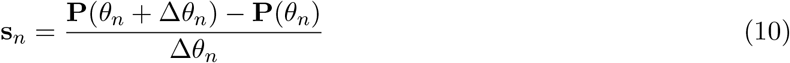

for some sufficiently small Δ*θ*_*n*_. Substituting the FSP approximation from (5) yields:

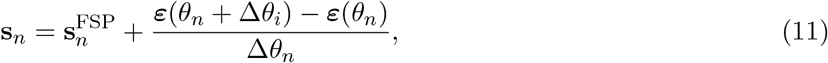

where 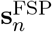 is estimated as the finite difference of the FSP solution, and ***ε***(*θ*_*n*_) and ***ε***(*θ*_*n*_ + Δ*θ*_*n*_) are the positive bounded errors from the FSP solutions (5).

Let *g* ≡ |**g**_**J**_|_1_ denote the total FSP error for the projection onto the states indexed by **J**. From (6), it is guaranteed that ***ε***(*θ*_*n*_) ≤ *g*_**J**_(*θ*_*n*_)**v**_1_ for some **v**_1_ ≥ 0 such that |**v**_1_|_1_ = 1 and ***ε***(*θ*_*n*_ + Δ*θ*_*n*_) ≤ *g*_**J**_(*θ* + Δ*θ*_*n*_)**v**_2_ for another **v**_2_ ≥ 0 such that |**v**_2_|_1_ = 1. Combining these, we can calculate FSP bounds on the error in the finite difference sensitivity function as:

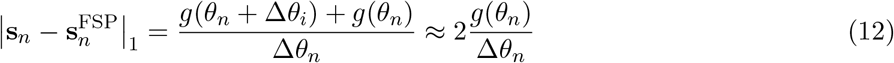

With this formulation the accuracy of the sensitivity approximation can be maintained by choosing the FSP projection **J** to ensure that *g*(*θ*_*n*_) ≪ Δ*θ*_*n*_.

### 2.6 FSP-Based Fisher Information Matrix

Now that we have approximated the sensitivity of the CME solution (12), we can compute the gradient of the log-likelihood function (7) to approximate the FIM:

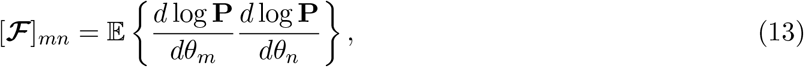

where the expectation is taken over the set of all possible independent data sets sampled from the CME distribution at specified times and experimental conditions. Since **P** = [*p*_*i*_] and [*dp*_*i*_/*dθ*_*m*_] = [**s**_*m*_]_*i*_, we can write the FIM as

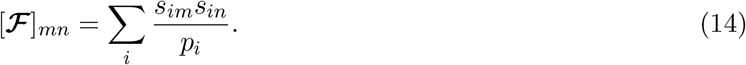

In matrix notation, this is written simply as:

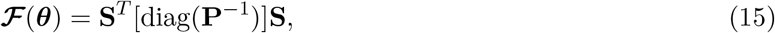

where **S** = [**s**_1_, …, **s**_*N*_] is the stoichiometry matrix, and [diag(**P**^−1^)] is a diagonal matrix with terms **P**^−1^. The FSP-FIM accuracy can be estimated numerically by propagating the estimated bounds on **P** and **S** in (6) and (12).

In the above formulation, **ℱ** is calculated for a single observation, at a single time, and for a single environmental condition. For independent snapshot experiments (e.g., ICC or smFISH), each cell is measured only once, and cell dynamics are assumed to be uncoupled from one another. If {*n*_*k*_} cells are chosen from a set of *K* experiments {*E*_1_, …, *E*_*K*_}, then the total FIM is the weighted sum:

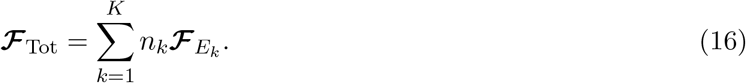

When estimating non-negative parameters with vastly different magnitudes, it is convenient to express parameters in log-space. Applying the relationship *du*/*d* log *θ* = (*du*/*dθ*)(*dθ*/*d* log *θ*) = (*du*/*dθ*)*θ* to (13) we can express **ℱ** in log-space as:

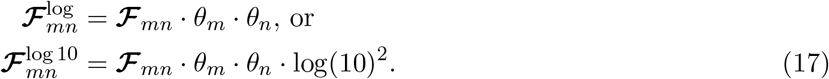

For the remainder of this paper, we will express parameter uncertainties and **ℱ** in log-10 space.

### 2.7 Cramér-Rao lower bound and experiment design

Recall that our goal is to estimate the parameters ***θ*** based on observations {**x**_*d*_, *t*_*d*_}. For a known model, the Cramér-Rao lower bound (CRLB) provides a (best-case) estimate for how much parameter uncertainty would result from different potential experiments. The CRLB states that for any unbiased estimator for ***θ***, the covariance of the estimate 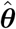 satisfies:

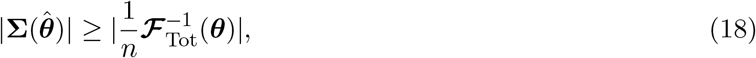

where **ℱ**_Tot_(***θ***) is from (15,16). Intuitively, the Fisher information matrix measures the amount of information that the observations contain about the parameter vector ***θ***.

To save effort or resources, one should choose experiments that will minimise some metric, termed the *utility function* of the estimator 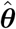. For the purposes of this study, we seek to minimize the volume of the expected parameter uncertainty for all (D-optimality, minimizing 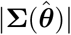) or a subset (G-optimality) of the model parameters. Other optimality criteria exist (e.g., E-optimality to minimise the largest eigenvalue, A-optimality to minimize the trace, etc.), and can be computed with minimal adjustment to the analyses below.

### 2.8 Bayesian Experiment Design

Experiment design can utilize either frequentist or Bayesian statistics. The latter is particularly attractive for sequential experiment design (SED), since although exact values of ***θ*** are unknown, one may have a prior distribution *P*(***θ***) or a posterior distribution computed from a previous round of parameter estimation, e.g., *P*(***θ***|𝒟) from (9), which can naturally serve as an input to the design of the next round. For D-optimality (in which we seek to minimize the |**Σ**(***θ***)|), we can compute a lower bound on the expected determinant of the estimator covariance as:

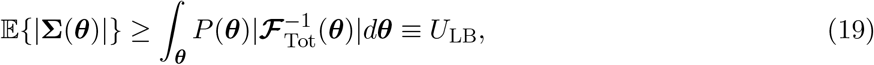

where the prior *P*(***θ***) is replaced with *P*(***θ***|D) in subsequent rounds. Although an analytical form for *P*(***θ***|𝒟) is not available, we can use the MHA from Sec. 2.4 to estimate:

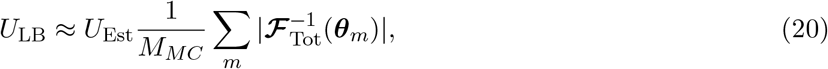

where {***θ***_*m*_} are independent Markov chain Monte-Carlo (MCMC) samples from *P*(***θ***|𝒟). With this estimate for the expected uncertainty, *U*_Est_, we can seek specific combinations of times and input signals to minimize this quantity. Suppose that we can perform *K* different combinations of time points and experimental conditions denoted by the the set of experiments {*E*_*k*_}. Let **n** = [*n*_1_, …, *n*_*K*_] denote an experiment design defining how many cells are to be collected for each possible experiment. If we restrict the experiment design to containing exactly *N*_*T*_ cells, then the optimal such design can be estimated:

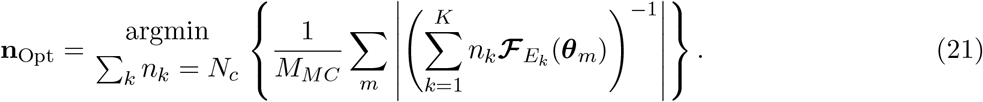

### 2.9 Glucocorticoid Receptor Localization Experiments

HeLa Kyoto cells were cultured in Dulbecco’s modified Eagle medium (DMEM, Thermo Fisher Scientific, 11,960–044) with 10% fetal bovine serum (FBS, Atlas Biologicals, F-0050-A), 10 U/mL penicillin/streptomycin (P/S, Thermo Fisher Scientific, 15140122), 1 mM L-glutamine (L-glut, Thermo Fisher Scientific, 25030081) in a humidified incubator at 37^°^C with 5% CO_2_. For immunocytochemistry (ICC), cells were seeded on cover glasses in DMEM supplemented with charcoal-stripped FBS (Sigma, F6765-500ML), P/S, and L-glutamine, 24 hours before the experiments. Dexamethasone (Dex, Sigma, D2915) was reconstituted in nuclease-free water to 63.7 mM and diluted to the required experimental concentrations; samples were removed from incubation only briefly for stimulant addition and mixing. The ICC protocol included fixing cells with 4% PFA (VWR, 100496-496), permeabilization with 0.1% Triton X-100 in PBS (Thermo Fisher, J66624-AE), blocking for nonspecific binding (Thermo Fisher, 01-6201), overnight incubation with a primary antibody (Abcam, EPR19621), followed by a secondary antibody (Thermo Fisher, A11008), 50 ng/mL DAPI (Thermo Fisher, D1306), and mounting for microscopy (Vector Laboratories, H-1000-10). Fluorescent images were acquired with an Olympus IX81 inverted spinning disk confocal (CSU22 head with quad dichroic and additional emission filter wheel to eliminate spectral crossover) microscope with 60x/1.42 NA oil immersion objective. Confocal z-stacks (0.5 *µ*m step-size, 27 stacks in each channel) were collected. ICC experiments used 2 high-power diode lasers with rapid (microsecond) switcher (405 nm DAPI (100 ms), 488 nM Alexa Fluor 488 (100 ms)).

### 2.10 Single-Cell Image Processing

To quantify ICC intensity signals, we implemented a Python image-processing pipeline [17]. First, we used the DAPI channel to segment the nuclei of all the cells in a given image using Cellpose [18]. After obtaining the nuclear masks, we created a pseudo-cytosol mask by expanding the nuclear mask using binary dilation. Then, the intensity was calculated for each cell using a maximum projection of the 3D microscope image and averaging the intensity values inside the nuclear and pseudo-cytosol mask independently. To avoid heterogeneity arising from cellular states, the top and bottom quartiles of cells (by nuclear size) were discarded to focus analyses on similarly sized cells. The GR fluorescence intensities were integrated over nuclear and cytoplasmic volume to record GR in arbitrary units of mass.

### 2.11 Sequential Experiment Design Algorithm

Given the above analysis tools, we can now specify a specific set of tasks for sequential experiment design (SED):

1. A set of data is collected according to the specified experiment design (i.e., collection and imaging of cells at specified time points and environmental conditions);
2. The FSP approach (Secs. 2.2 and 2.3) is used to maximize the likelihood of the resulting data (7);
3. Metropolis-Hastings sampling (2.4) is conducted to collect *N*_*MH*_ samples and estimate the posterior distribution (9) given the new data;
4. The *M*_MC_ sets of FIMs are calculated (Sec. 2.6) using parameter samples selected uniformly from the MHA chain and for all possible experiment designs; and
5. The *N*_*c*_ cells of the next experiment are optimally allocated to minimize the expected uncertainty volume (20), and the next round of experiments is performed.

Our goal is to minimize the uncertainty volume, given by the determinant, |**Σ**(***θ***|. To estimate performance for future experiments, we sample the posterior of the previous stage and compute the average inverse determinant of the FIM using (20). For simulated data, where a ground-truth model is available, we also compute the inverse FIM determinant for the true parameters. To quantify performance of past experiments, we use MHA samples to estimate the determinant of the posterior parameter covariance matrix.

We consider two SED strategies: The *Bayesian FIM-Based Design*(**BFBD**) chooses the optimal allocation of signals |*s*|_*k*_ ∈ 𝒮 and measurement times *t*_*k*_ ∈ 𝒯 for each of *k* = 1, 2, … groups of *N*_*c*_ new cells at each stage. The *Random Design*(**RD**) chooses the input and measurement times at random. Other strategies are certainly possible given the tools described above, but these are left for future investigations.

All analyses are conducted in Matlab using the Stochastic System Identification Toolkit; scripts are available at: [19].

## 3 Results

To demonstrate the SED approach, we consider three models of varying complexity. For each model, we start with a rough guess for the parameters and a preliminary data set (simulated data in the first two examples; experimental data in the third). We then execute the algorithm described in Sec. 2.11 and compare the results for different ED strategies.

### 3.1 Example 1 - Transcription with Modulated Rate

In the first example, we consider a simple two-reaction transcription model. RNA production is modeled as

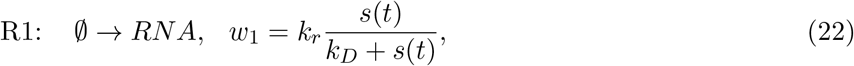

where ∅ denotes an independence from reactants (i.e., a zeroth order reaction), *k*_*r*_ is the fully active transcription rate, and *s*(*t*) = |*s*|*u*(*t*), is a step-input induction signal of magnitude |*s*| that activates the promoter through a Michaelis-Menten interaction with dissociation constant *k*_*D*_. Degradation is modeled as standard linear decay:

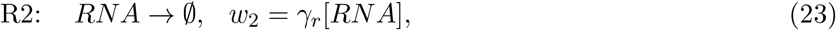

where *γ*_*r*_ is the degradation rate. To generate simulated data, we choose the parameters: *k*_*r*_ = 10 min^−1^, *k*_*D*_ = 5 nM, and *γ*_*r*_ = 0.3 min^−1^. The system is assumed to begin with an initial inactivated condition of zero RNA at *t* = 0.

The SED begins with a first experiment with a step magnitude of |*s*| = 1 nM, and 15 cells are measured at each time {0, 4, 6.5, 10} min. The set of allowable subsequent experiments at each stage is any combination of 60 new cells under signal magnitude |*s*|_*i*_ ∈ 𝒮 = {1, 2, …, 10} nM and collected at times *t*_*i*_ ∈ 𝒯 = {0, 0.5, 1, …, 10} min. We consider eight experiment rounds, including the initial experiment, leading to a final set of 8×60=480 total cells. The fitting routine has access to the number of RNA in each cell, as needed to compute the likelihood function (7).

We seek to minimize the determinant of the log10 covariance matrix for parameters. Before fitting, we assume a log-normal prior for the distributions of {*k*_r_, *γ*_r_, *k*_D_} with log10-mean of zero and a log10-standard deviation of one, corresponding to a starting uncertainty of |**Σ**|_Prior_ = 1.

Figure 1B shows the convergence of the uncertainty over stages two through eight of the sequential experiments for both the **BFBD** (red) and **RD** strategies (black). For each strategy, three computations are shown. The dashed lines correspond to the uncertainty volume predicted by the FSI-FIM using the parameter estimates from the previous round of experiments, and the solid line shows the actual uncertainty volume computed from the MHA for the current data set. The square markers show the uncertainty volume that one would expect for that experiment if the true parameters were known in advance. For all rounds, we find that the MHA algorithm predicts a posterior uncertainty that is in excellent agreement with what would have been expected if the true parameters were known (compare solid lines with square markers of the same color). For the third and subsequent round of experiments, i.e., after 120 cells are included in the data set, we find that the FIM prediction of the uncertainty based on the current parameters is also in excellent agreement with the true values and with the MHA estimates (compare dashed lines with square markers and solid lines of the same color).

**Figure 1:**
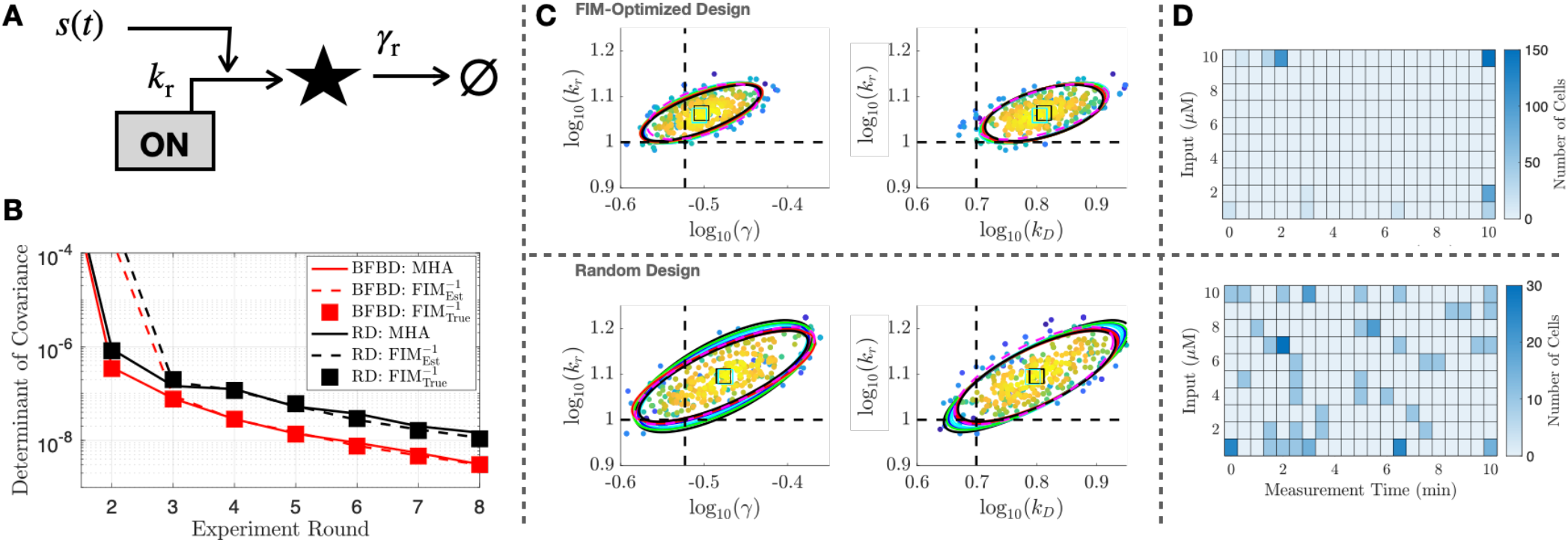
Poisson Transcription with Modulated Production Rate. (**A**) Schematic of model (see text for propensity functions and parameters). (**B**) Determinant of log10-covariance matrix, versus experiment round for **BFBD** (red) and **RD** (black). Uncertainty is determined from Metropolis-Hastings sampling (solid lines), predicted using FIM with parameters estimated from previous round (dashed); or based on FIM with exact parameters (squares). (**C**) Joint parameter uncertainties in experiment round 4 for BFBD (top) and **RD** (bottom) designs. MHA samples are shown with dots; MHA 90% CI is shown in dashed ellipse; 10 FIM predictions (using parameter samples from round 3) are shown in solid ellipses. True parameter values are denoted with dashed horizontal and vertical lines. (**D**) Number of cells sampled at each input concentration and at each time after eight rounds of experiments. Both experiment design methods consider 60 new cells in each round.

The agreement between the FIM-estimated uncertainty and the MHA-based sampling of the posterior is also illustrated in Fig. 1C, which compares the joint parameter uncertainty estimates based on FIM predictions after three rounds of experiments (solid ellipses) with the actual MHA uncertainty estimates (dots and dashed ellipse) computed after collecting and fitting new data in the fourth round. The true parameters are denoted by vertical and horizontal dashed lines.

Comparing the results for the **BFBD** and **RD** designs, we find that the **BFBD** reduces uncertainty. For example, the uncertainty volume achieved by **RD** after eight rounds is found to be |**Σ**_MHA_|_RD:r8_ = 1.46 × 10^−8^, whereas **BFBD** achieves a slightly better performance of |**Σ**_MHA_|_BDBD:r5_ = 1.39 × 10^−8^ in only five rounds.

Finally, Fig. 1D shows the number of cells measured at each time point and for each input level for the two different strategies. We find that the optimized ED is much simpler than what one might intuit. Specifically, the optimized sequential experiments prioritize cell collection at the highest induction level (|*s*| = 10 nM), where 150 cells are measured at early times (10, 30, and 110 cells at 0.5, 1.5 and 2.5 min, respectively) and 150 cells are measured at the final time (10 min). Of the remaining 120 cells in rounds 2-8, 35 are collected at the final time (10 min) and lowest induction concentration (|*s*| = 1 nM); and 100 are collected at the second lowest induction level (|*s*| = 2 nM with 10 at time *t* = 3 min and 90 at *t* = 10 min).

### 3.2 Example 2 - Bursting Gene Expression Model with Unknown Induction Mechanism

We next consider a bursting transcription model, where a promoter can switch between two potential configurations: an ON state with transcription at a constant rate *k*_r_, and a silent OFF state. The model (see Fig. 2A) consists of four potential reactions defined as:

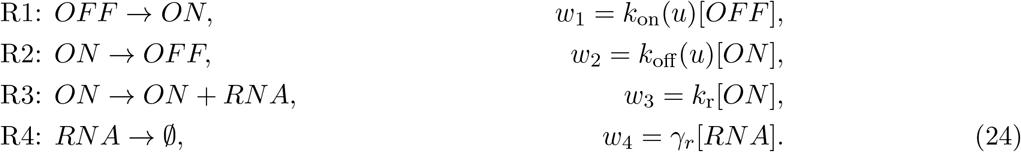

**Figure 2:**
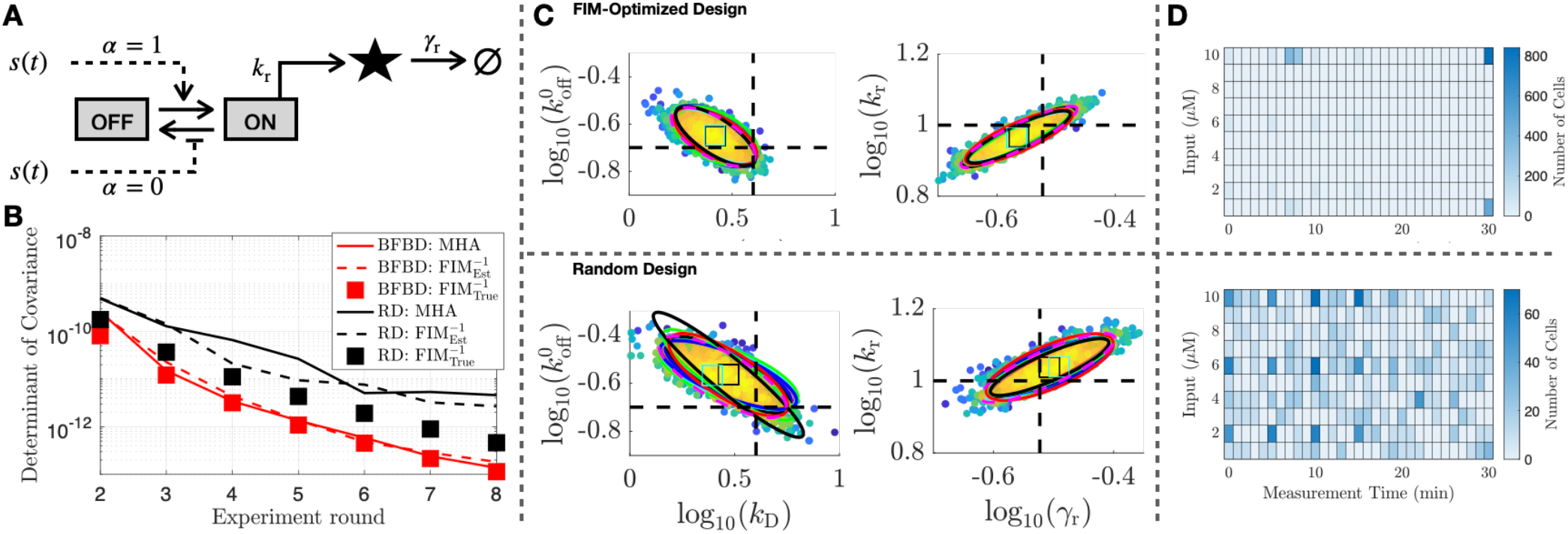
Bursting Gene Expression Model with Uncertain Induction Mechanism. (**A**) Schematic of model consisting of two states, ‘ON’ and ‘OFF’. Reactions between these states can depend on an inducer concentration through one of two potential mechanisms: burst frequency control (*α* = 1) and burst size control (*α* = 0). See text for specific propensity functions and parameters. (**B-D**) Same as in Fig. 1, but where the model now has six parameters (only two of 15 possible pairs are shown). Both experiment design methods consider 300 new cells in each round.

The rates of switching between the two states depend on the external environment according to:

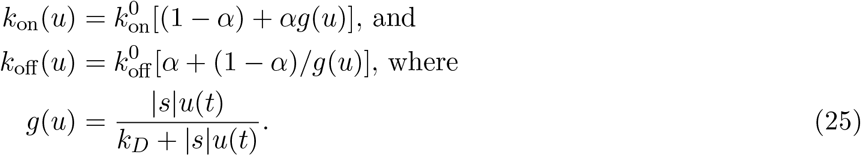

This specific form of the inputs is such that when *α* = 1, the control signal modulates the rate of activation, and when *α* = 0, the input signal modulates the duration of activity. As in the previous example, we assume that step inputs can be given to the system at time *t* = 0 with variable magnitudes |*s*|. Note that the average steady-state gene expression is

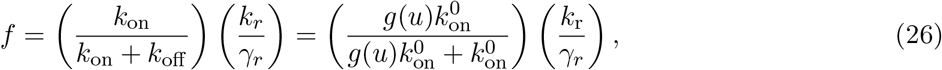

which is not a function of *α*, and the mechanism of action cannot be determined from average steady-state expression.

To generate simulated data for subsequent fitting and parameter identification, the ground-truth parameters of the model have been set to:

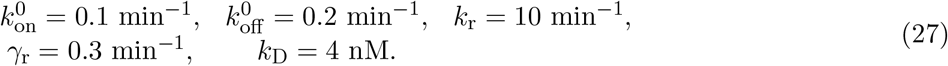

The true value of *α* was set to zero, corresponding to modulation of the gene deactivation rate.

Before fitting the first round of data, we assume a log-normal prior for the distributions of parameters 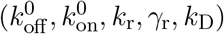 to have log10-means of zero and log10-standard deviations of two corresponding to a starting uncertainty volume for these parameters of |**Σ**|_Prior_ = (2^2^)^5^ = 1024. Because the mechanism parameter *α* is constrained to lie between zero (burst-length control) and one (burst-frequency control), whereas the remaining parameters can take on any positive quantity, we introduce the transformation *a* = tan(*απ*/2) before fitting. For the initial stage, we assume that *a* also has a log-normal prior with a log10-mean of zero and a very broad log10-standard deviation of four. Furthermore, because the transformed variable *a* is now expected to approach either zero (burst-length control) or infinity (burst-frequency control), its log-sensitivity is expected to approach zero when the correct mechanism has been identified. Therefore, in the SED, we fit and quantify uncertainty in all parameters, but use G-optimality to minimize the determinant of the log10 covariance matrix for all parameters except *a*.

The SED begins with three experiments with step input magnitudes of |*s*|_1_ ∈ {2, 6, 10} nM. For each experiment, 50 cells are measured at each time {0, 5, 10, 15} min. For each subsequent experiment, we collect a set of 300 new cells (in groups of 10 cells) under any signal magnitude |*s*|_*i*_ ∈ 𝒮 = {1, 2, …, 10} nM and any times *t*_*i*_ ∈ 𝒯 = {0, 1, …, 30} min. We consider eight experiment rounds, including the initial experiment, for a final set of 600+7×300=2700 total cells. The fitting routine has access to the number of RNA in each cell, but the gene state is treated as unobservable.

Figure 2B shows the uncertainty versus the experiment round for the **BFBD** (red) and **RD** (black) strategies for the MHA sampled posterior (solid lines), the FIM uncertainty predictions from the previous round (dashed lines), and the FIM lower bound using the exact parameters (squares). As before, the estimated and exact FIM are in excellent agreement for all cases (compare dashed lines and squares of same color). For **BFBD**, these also agree strongly with the posterior uncertainty (compare all red lines and points).

Figure 2C shows the joint uncertainties for parameters pairs 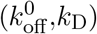 and 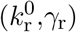 after four experiment rounds. The **BFBD** strategy leads to smaller parameter uncertainties (Fig. 2B, compare red and black lines), especially for 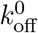 and *k*_D_ (Fig. 2C Left, compare top and bottom). Furthermore, Fig. 2 shows that the **BFBD** achieves its higher performance despite a very simple experiment design in which all new cells are measured at the highest (|*s*| = 10 nM, 1510 out of 2400 cells) or the lowest ((|*s*| = 1 nM, 590 out of 2400 cells) induction level and only at a handful of times at the beginning or very end of the time course. Both strategies successfully identify the regulatory mechanism after just a few rounds, correctly setting *a* ≪ 1, i.e. burst-frequency modulation.

### 3.3 Example 3 - Induction of Glucorticoid Receptor Cytoplasm-to-Nucleus Transport

For a third demonstration of SED strategies, we consider a model of Dexamethasone stimulation of the transport of the glucocorticoid receptor (GR) from the cytoplasm to the nucleus. In this simple model (Fig. 3A), we consider two species, labeled as *GR*_cyt_ when the receptor is in the cytoplasm and *GR*_nuc_ when it is in the nucleus. Production of GR is assumed to occur only in the cytoplasm, and degradation is assumed to occur only in the nucleus. Thus, there are four reactions:

**Figure 3:**
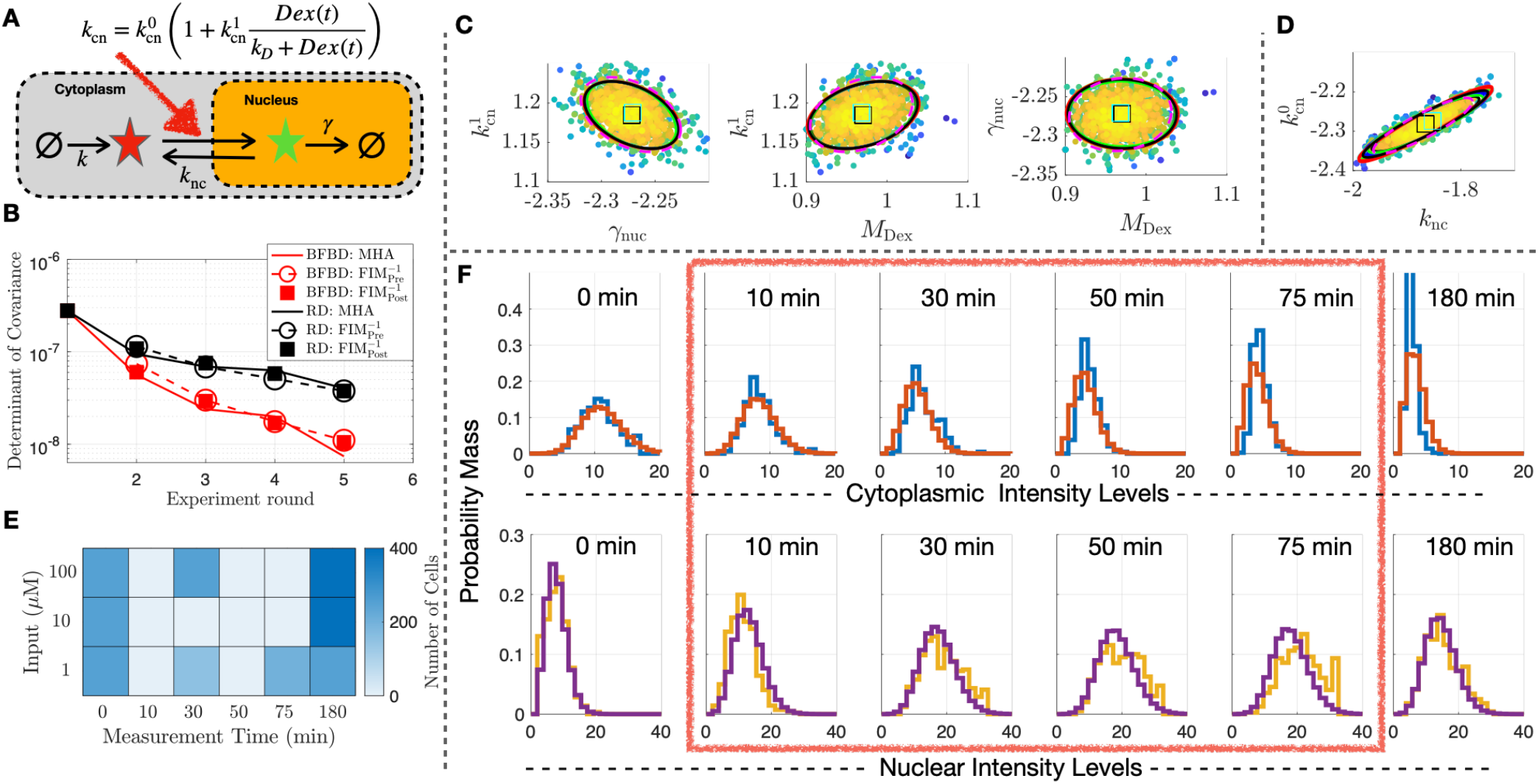
GR Localization. (**A**) Model Schematic: GR translocates between cytoplasm and nucleus depending on Dex concentration, production occurs in the cytoplasm, degradation occurs in the nucleus. (**B**) Uncertainty volume for chosen parameters versus iteration. (**C**) Joint uncertainties for parameters 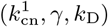 as estimated with FIM before the 4th experiment and using MHA after the 4th experiment. (**D**) Uncertainties for 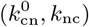. (**E**) Collective experimental design after five rounds. Only 9 out of 18 combinations were chosen for data collection. (**F**) Model predictions for the distributions of cytoplasmic (top) and nuclear (bottom) concentration for Dex = 10 nM. Data in the red box were *never* selected for use in the parameter estimation.

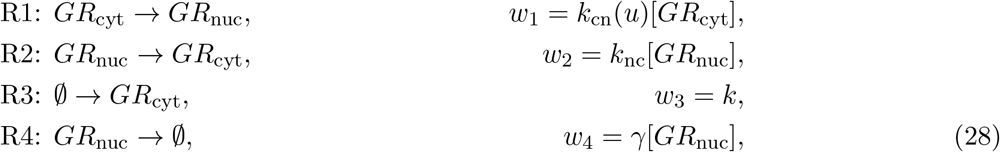

In R1, the cytoplasm-to-nucleus transport rate is defined:

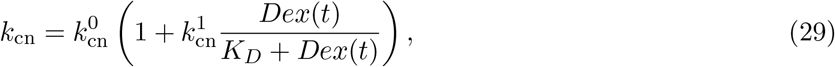

where *Dex*(*t*) is added as a step input at time *t* = 0.

As in the previous two examples, our goal is to identify the parameters of the model. To achieve this, we have quantified the GR in each location (at single-cell resolution) using immunocytochemistry at induction levels of {1, 10, 100} nM Dex, and at time points of {0, 10, 30, 50, 75, 180} min following stimulation. For each time point, we measured GR localization intensities for between 284 and 473 cells, after removing the cells with the 25% largest and 25% smallest nuclei in efforts to reduce effects of extrinsic noise. Brief details are included in Section II.J-K, but full details of the experimental approach and a more thorough investigation of a model for the GR dynamics and its downstream effects on gene expression will be described elsewhere. For the purpose of this study, we wish to know: (1) if we could have achieved equivalent understanding through a simpler sequence of experiments, and (2) what experiments should we do next to achieve improved identification of the model.

Drawing inspiration from the previous examples where optimal experiments prioritized high induction levels and the late time points, we started with initial experiments of 150 cells each at times *t* = {0, 30, 180} min at {1, 100} nM Dex. We then perform six rounds of SED using the **BFBD** approach and add 300 new cells in each iteration. Although all six parameters 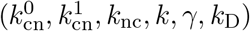 are treated as unknown, we use *G*-optimality criteria in which we seek only to minimize the uncertainty volume for only 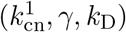.

Because there is no known true model in this case, at each stage we compare the MHA results to a FIM-based prediction from the previous round (circles and dashed lines) and to an updated FIM-based estimate based on the new parameters in the current round (squares). From Figure 3B, it can be seen that the MHA and FIM results are in excellent agreement, and also that the **BFBD** performs better than the **RD**. Figure 3C shows the MHA and FIM-predicted uncertainty ellipses for the three parameters used in the optimality criteria, and Fig. 3D shows the same for two of the remaining parameters. In all cases the FIM provides an excellent prediction for the uncertainty, which is encouraging considering that the FIM calculation is a necessary step for creating an efficient MHA analysis and takes orders of magnitude less time to complete.

Figure 3E shows the collective design combining all six rounds of experiments. We find that the Bayesian-FIM approach led to a far simpler experiment than one would have designed by chance (and indeed much simpler than the intuitively-designed experiment we had originally conducted). In particular, the SED only selected initial and final time-points from the 10 nM induction experiment. As a final validation of the approach, Fig. 3 shows that the identified model accurately predicts the response distributions for these left-out time points. This fact that we can make accurate predictions for all left-out data sets further confirms that these time points were not informative and therefore not needed for parameter inference.

The parameters (mean ± SD) determined in the fitting procedure after 6 rounds were: 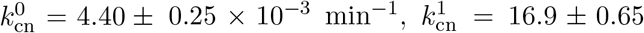, *k*_nc_ = 0.0104 ± 0.00092 min^−1^, *k* = 0.0149 ± 0.00079 min^−1^, *γ* = 5.71 ± 0.23 × 10^−3^ min^−1^, *k*_D_ = 10.78 ± 0.53 nM.

## 4 CONCLUSIONS

We considered sequential experiment design to identify discrete stochastic models to predict single-cell responses. In our procedure, small datasets are collected, stochastic models are defined using the CME framework, parameter uncertainties are found via Bayesian sampling, and the FIM is used to suggest the next experiment round. We validated these methods on three different models (two simulated and one experimental), each subject to an external signal. We found that FSP-FIM consistently made accurate predictions for the posterior uncertainties and that sequential FIM-based designs led to better identification with less required data.

These promising preliminary results provide motivation to expand the applicability of model-guided experiment design. Although we considered only piece-wise constant inputs, our analyses generalize to arbitrary time-varying inputs *s*(*t*). More diverse signals could enable identification from even less data [20]. Although we assumed ideal data without measurement errors; it will be interesting to see how experiment designs change under realistic distortions due to microscopy resolution, labeling inefficiencies, or image-processing errors, which can be incorporated into the FIM analyses [13]. In the GR example, we assumed a mechanism of Dex action and only consider the translocation process, but it would be interesting to design experiments to determine this mechanism or how GR affects downstream transcription.

Here, we consider only snapshot data where every measurement is independent, as collected using common endogenous post-fixation, single-cell experiments such as smFISH, ICC, flow cytometry, or sequencing. However, live cell trajectory measurements can be collected using genetic manipulations to include fluorescent reporters or through labeling with fragmented antibodies (e.g., see [21]). For such systems, the FSP tools above can be adapted to compute the FIM for trajectory measurements [22], or one could approximate the model as a stochastic differential equation and apply filtering methods [23]. The presented FSP-based approach requires integration of linear ODEs whose dimensions scale exponentially with the number of chemical species included in the model. While some aspects of this issue can be addressed through model reductions (e.g., based on time scale separations [24], coarse grained models [25], and balanced truncations [26]), many CME models remain intractable, and further model reduction approaches are needed. Moreover, even when the CME can be solved efficiently, if the model includes large numbers of parameters, it may not be possible to effectively search parameter space or collect sufficient data to determine those parameters. Given the potential benefits of model-guided experiment design for those cases where the FIM can be calculated directly as shown here, there remains a strong motivation for future computational and theoretical developments to address or circumvent these limitations.

